# Evolution of a *cis*-acting SNP that controls Type VI Secretion in *Vibrio cholerae*

**DOI:** 10.1101/2022.01.11.475911

**Authors:** Siu Lung Ng, Sophia Kammann, Gabi Steinbach, Tobias Hoffmann, Peter J Yunker, Brian K. Hammer

## Abstract

Mutations in regulatory mechanisms that control gene expression contribute to phenotypic diversity and thus facilitate the adaptation of microbes and other organisms to new niches. Comparative genomics can be used to infer rewiring of regulatory architecture based on large effect mutations like loss or acquisition of transcription factors but may be insufficient to identify small changes in non-coding, intergenic DNA sequence of regulatory elements that drive phenotypic divergence. In human-derived *Vibrio cholerae*, the response to distinct chemical cues triggers production of multiple transcription factors that can regulate the Type VI Secretion System (T6), a broadly distributed weapon for interbacterial competition. However, to date, the signaling network remains poorly understood because no regulatory element has been identified for the major T6 locus. Here we identify a conserved *cis*-acting single nucleotide polymorphism (SNP) controlling T6 transcription and activity. Sequence alignment of the T6 regulatory region from diverse *V. cholerae* strains revealed conservation of the SNP that we rewired to interconvert *V. cholerae* T6 activity between chitin-inducible and constitutive states. This study supports a model of pathogen evolution through a non-coding *cis*-regulatory mutation and preexisting, active transcription factors that confers a different fitness advantage to tightly regulated strains inside a human host and unfettered strains adapted to environmental niches.

**Importance:** Organisms sense external cues with regulatory circuits that trigger the production of transcription factors, which bind specific DNA sequences at promoters (“*cis*” regulatory elements) to activate target genes. Mutations of transcription factors or their regulatory elements create phenotypic diversity, allowing exploitation of new niches. Waterborne pathogen *Vibrio cholerae* encodes the Type VI Secretion System “nanoweapon” to kill competitor cells when activated. Despite identification of several transcription factors, no regulatory element has been identified in the promoter of the major Type VI locus, to date. Combining phenotypic, genetic, and genomic analysis of diverse *V. cholerae* strains, we discovered a single nucleotide polymorphism in the Type VI promoter that switches its killing activity between a constitutive state beneficial outside hosts and an inducible state for constraint in a host. Our results support a role for non-coding DNA in adaptation of this pathogen.

## Introduction

A central role in the dynamic, temporal control of gene expression is played by transcription factors (TFs), diffusible “*trans”* products that bind to molecular switches within DNA sequences termed “*cis”*-regulatory elements (CREs). In eukaryotes, which lack horizontal gene transfer (HGT), mutations in CREs that alter TF binding sites are major contributors to phenotypic diversity (1-3). In bacteria, pervasive HGT of TFs can alter entire regulatory circuits that allow adaptation to new niches, as prominently demonstrated in *Vibrio fischeri*, where host range is altered by the presence or absence of RcsS, a TF of biofilm and colonization genes (4, 5). By contrast, specific mutations at CREs in non-coding DNA are more difficult to identify and receive less attention as drivers of phenotypic divergence and evolutionary adaptation (6). Thus, elucidation of how microbes adapt to new niches, a process of fundamental importance in bacterial pathogenesis, requires coupling of genome-wide computational methods with experimental approaches to map the *cis-* and *trans*-regulatory interactions across and within species.

To understand how mutations play a role in microbial adaptation, pathogenic viruses and bacteria with lifestyles that exploit niches within and outside a human host are of great interest. Following ingestion, pandemic strains of the bacterium *Vibrio cholerae* can colonize the human gastrointestinal tract and secrete the cholera toxin that leads to the often fatal diarrhea responsible for seven pandemics to date (7-9). Conversely, *V. cholerae* isolated from non-human niches lack the horizontally-acquired prophage that carries the cholera toxin, and cause mild illness (10). By contrast, all sequenced *V. cholerae* encode a Type VI Secretion System (T6), a broadly distributed “nano-harpoon” weapon that injects toxic effector proteins into neighboring bacterial cells, leading to cell envelope damage and cell lysis (11, 12). Due to its broad distribution among bacteria including those of the human gut, there is intense interest in understanding the T6 interactions between our microbiota and foreign pathogens, and whether they can be manipulated to influence health (13).

*V. cholerae* obtained from humans carry a limited arsenal of effectors and a T6 believed to be tailored for *in vivo* success (11, 14-19), while strains from non-human niches encode a more diverse effector repertoire (11, 14, 20, 21). To date, however, adaptative evolution mechanisms of T6 regulation in *V. cholerae* derived from non-human sources have largely been overlooked. Studies of human-derived strains identify two primary TFs for T6 activation (22-26). T6 control in pandemic strains (e.g. C6706 and A1552) requires QstR (Quorum-Sensing and Chitin-Dependent Regulator), which integrates multiple external cues (27-30), and contains a DNA binding domain postulated to interact with a presumptive CRE of the major T6 gene cluster (23, 27). T6 regulation in non-pandemic strain V52, which causes mild disease, requires TfoY, modulatable by intracellular signals, including cyclic di-GMP (25, 26). How QstR and TfoY control T6 transcription remains elusive, with no T6 CRE yet described. Elucidation of the differences in intraspecies T6 regulatory mechanisms between diverse *V. cholerae* isolates will provide insights into how pathogens emerge from nonpathogenic progenitors.

To understand the regulatory differences in *V. cholerae* strains, we examine here several environmental isolates that exhibit T6-mediated killing (31). Despite encoding functional signaling circuity and TFs, we find that QstR is dispensable for killing and that TfoY plays only a minor role in the strains tested. Thus, existing regulatory models fail to explain the T6 control in *V. cholerae* from human and non-human sources. Genomic analysis identifies one conserved non-coding single-nucleotide polymorphism (SNP) that we show interconverts *V. cholerae* T6 activity between chitin-inducible and constitutive states, which are QstR-dependent and TfoY-independent, respectively. We demonstrate that non-coding SNPs can rewire *cis*-regulatory elements to aid in adaptation of bacteria to different niches, including the human host.

## Results and Discussion

### Constitutive, *in vitro* T6 activity requires neither QstR nor TfoY

In pandemic C6706, high cell density conditions (HCD) and chitin are required for induction of *qstR* which leads to activation of T6 genes. In the absence of chitin, C6706 with *qstR* expressed from a heterologous promoter (defined here as *qstR*\*) reduces survival of *Escherichia coli* “target” cells in co-culture by over 4-orders of magnitude (∼10,000), compared to wildtype (WT) C6706, a T6^-^ strain with a mutation in an essential structural gene (*ΔvasK*), and a strain with a *ΔqstR* mutation (Fig. 1A) (29). Deletion of *tfoY* does not reduce the killing activity of the T6^+^ *qstR*\* strain, but eliminates the robust killing in the non-pandemic strain V52 (serogroup O37), which requires TfoY but not QstR (Fig. 1B) (26).

**Figure 1.**
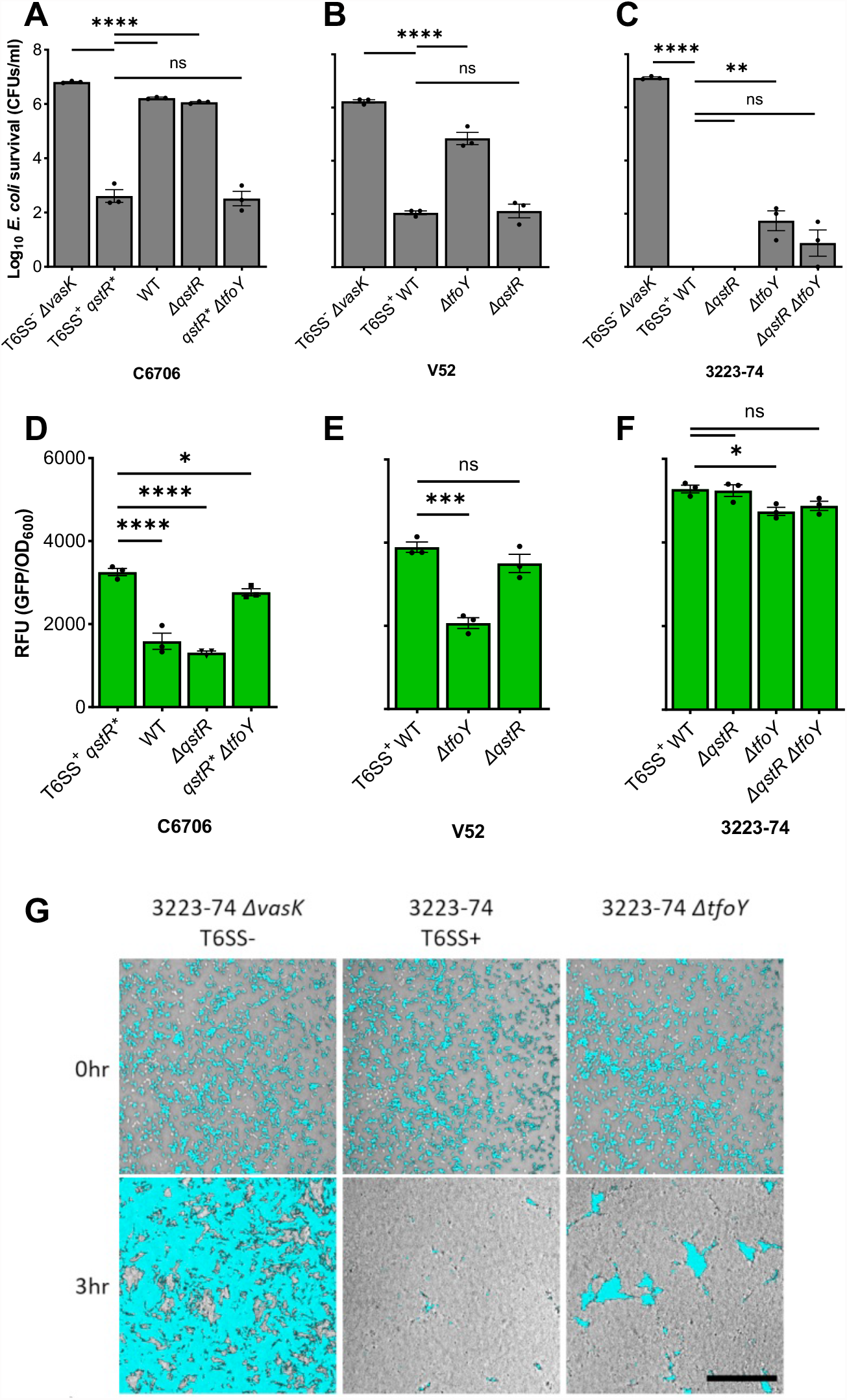
*Vibrio cholerae* 3223-74 T6 activity is QstR- and TfoY-independent. (A-C) *V. cholerae* strains with the indicated genotypes were co-cultured with chloramphenicol resistant (Cm^r^) *E. coli* followed by determination of *E. coli* survival by counting of colony forming units (CFUs) on LB agar with Cm. (D-F) Fluorescence levels are from reporters with *gfp* fused to the intergenic region 5’ of *vipA* derived from the three strains shown. The mean value ± S.E. of three independent co-cultures (A-C) and monocultures (D-F) are shown from one experiment, with similar results obtained in at least two other independent experiments. A one-way ANOVA with Dunnett post-hoc test was conducted to determine the significance: ns denotes not significant, ****p ≤ 0.0001, ***p ≤ 0.001, **p ≤ 0.01, *p ≤ 0.05. (G) *E. coli* cells expressing constitutive *gfp* were competed against 3223-74, with the same frame imaged at 0 h and 3 h by confocal microscopy. In the images, *gfp* signal from the *E. coli* is overlaid on top of bright-light images of the co-culture. Scale bar = 50 µm.

To determine whether QstR or TfoY participates in control of the T6 in non-human strains, we examined 3223-74, a genetically-amenable, T6-proficient environmental strain (31). Like V52, 3223-74 does not require QstR to efficiently kill *E. coli* in conditions without chitin, but surprisingly, also does not require TfoY. Isogenic strains carrying the *ΔtfoY* and *ΔqstR ΔtfoY* mutations retain >99.99% killing activity, with only modest *E. coli* survival (Fig. 1C). Gene fusions of the 5’ intergenic region (IGR) of the major T6 cluster of each strain fused to green fluorescent protein (*gfp*) confirm that transcriptional differences account for the killing observed, with maximal *gfp* expression mirroring activity (i.e. low *E. coli* survival with high *gfp* expression, and *vice versa*) (Fig. 1D-F). Confocal microscopy reinforces the negligible role of TfoY on killing by 3223-74, with a Δ*tfoY* mutation having little effect on killing WT (Fig. 1G). Transcription of plasmid-borne reporters is significantly higher in *V. cholerae* than in *E. coli* (Fig. S1), supporting a hypothesis that an additional *V. cholerae*-specific regulator of the T6 may remain to be identified (Fig. S1).

To probe each strain’s T6-related regulatory circuitry, we measured canonical behaviors under control of HapR, QstR and TfoY; quorum sensing (QS) controlled bioluminescence, natural transformation, and motility, respectively (32-34). As expected, each TF is intact in C6706; but like several *V. cholerae* strains, V52 lacks a functional *hapR* gene that prevents QS and natural transformation (35, 36). Nonetheless, V52 encodes a functional *tfoY* that controls motility (Fig. 2A-B) (37). Interestingly, the regulatory circuity of *V. cholerae* 3223-74 is intact, like C6706, confirming that it encodes functional TFs (Fig. 2C), which are nonetheless expendable for T6-mediated killing. Because transposon mutagenesis failed to identify a novel T6 activator (not shown), we suspect regulation may be complex, perhaps involving more than one TF specific to *V. cholerae*. Nucleoid Associated Proteins (NAPs) that bind DNA both specifically and non-specifically (38) may also contribute, since they are present in both species, likely regulated differently (39), and participate in regulation of many promoters in numerous bacteria including *Vibrios* (40).

**Figure 2.**
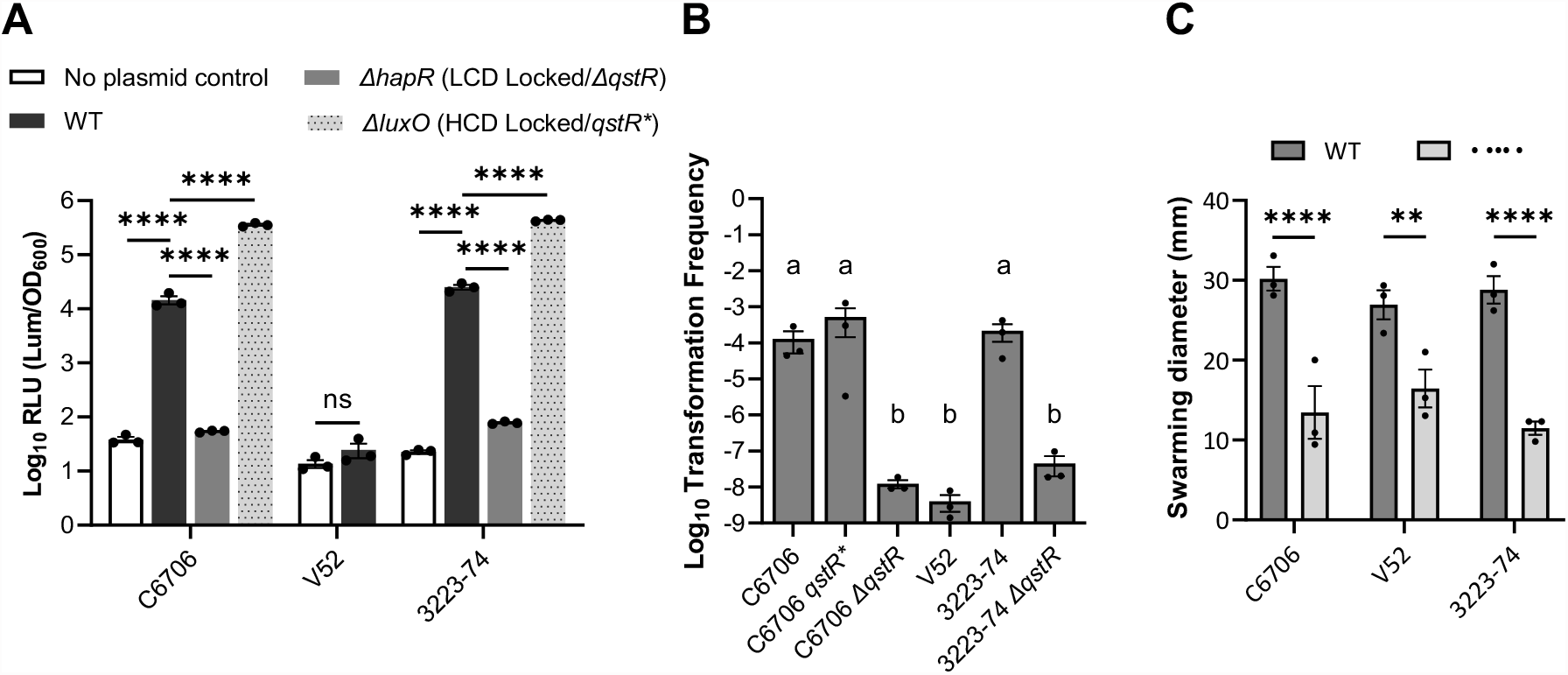
*Vibrio cholerae* 3223-74 encodes functional HapR, QstR, and TfoY. (A) *V. cholerae* strains were grown in liquid LB with relative luminescence units per OD_600_ measured at HCD (OD_600_ = 0.6-0.8). Statistical analyses were conducted with one-way ANOVA with Tukey post-hoc test (C6706 and 3223-74) and one-tailed student’s t-test (V52). LCD – Low Cell Density. (B) *V. cholerae* strains with the indicated genotypes were grown in ASW with crab shell and exogenous Spec-marked genomic DNA. Transformation frequency = Spec^r^ CFU ml^−1^ / total CFU ml^−1^. Statistical analyses were conducted with one-way ANOVA with Tukey post-hoc test. Significance is denoted by letters. (C) *V. cholerae* strains were inoculated on 0.3% LB agar and grew overnight. Statistical analyses were conducted with one-tailed student’s t-test. Colony diameters were physically measured from the furthest edges. All data shown are the mean ± S.E. from one experiment, with similar results were obtained in at least two other independent experiments. ns: not significant, ****p ≤ 0.0001, **p ≤ 0.01.

### A SNP in the T6 intergenic region confers QstR-dependency

Human and environmental isolates of *V. cholerae* we have characterized prior (31) share ≥97% average nucleotide identity with many chromosomal differences (11), but inspection of the T6 IGRs of C6706, V52 and 3223-74 revealed only 17 SNPs and 3 multinucleotide polymorphisms (Fig. 3A), which we hypothesized could contribute to the differences in T6 transcription and killing activity observed. To address this, we replaced the T6 IGR of C6706 on the chromosome with that from V52 and 3223-74 and measured killing activity. While C6706 carrying the *qstR\** allele, but not WT, adeptly kills *E. coli*, both IGR replacements increase the killing efficiency of WT C6706 by 5- to 6-orders of magnitude (Fig. 3B), mimicking the robust killing observed by WT V52 and 3223-74 (Fig. 1B-C). Deletion of *tfoY* in C6706 with V52’s IGR increases *E. coli* survival (∼ 2-logs), as observed with V52, but does not alter *E. coli* survival with 3223-74’s IGR (Fig. 3B). Chromosomal transcriptional *gfp* reporters with identical mutations were elevated relative to WT C6706 in each IGR replacement strain (Fig. 3C), consistent with the enhanced killing detected. These results support a hypothesis that a novel CRE lies within the IGR 5’ of the T6 locus, despite a lack of any known direct TF-DNA interactions at this locus identified to date.

**Figure 3.**
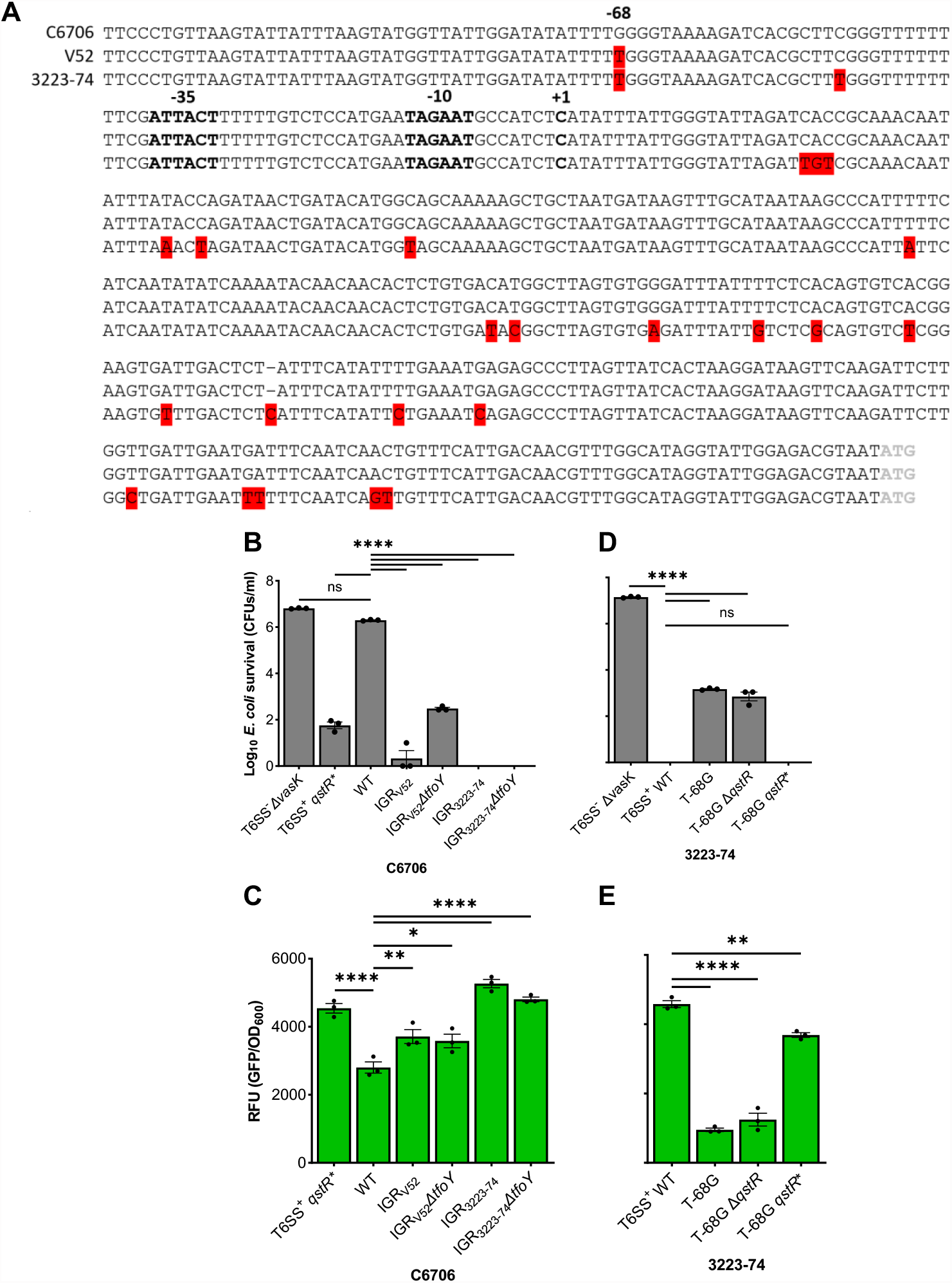
G-388T mutation abolishes QstR dependence in C6706 and T-388G confers QstR dependence to 3223-74. (A) Alignment of the IGR upstream of *vipA* was conducted using MUSCLE. SNPs and MNPs are highlighted in red, one gap indicated with a “–”, the putative promoter and the transcriptional start site (TSS; +1) in bold, and the start codon of *vipA* in grey. (B) the C6706 5’ IGR of *vipA* was replaced with the IGR from either V52 or 3223-74. (D) A T-68G mutation in the 5’ IGR of *vipA* was introduced into 3223-74 with different *qstR* alleles. Competition assays were conducted by co-culturing *V. cholerae* killers and Cm^r^ *E. coli* target followed by determination of *E. coli* survival by counting of colony forming units (CFUs) on LB agar with Cm. (C, E) Shown are fluorescence levels of transcriptional reporters with *gfp* fused to corresponding IGRs of *vipA* expressed in either C6706 (C) or 3223-74 (E). The mean value ± S.E. of three independent co-cultures (B and D) and monocultures (C and E) are shown from one experiment, with similar results were obtained in at least two other independent experiments. A one-way ANOVA with Dunnett post-hoc test was conducted to determine the significance - ns: not significant, ****p ≤ 0.0001, **p ≤ 0.01, *p ≤ 0.05.

To begin mapping the T6 IGR region and SNP locations, we experimentally determined the transcriptional start site (+1) by 5’ Rapid Amplification of cDNA Ends (Methods). The +1 of transcription resides 320 nucleotides (nt) 5’ of the ATG of the first T6 gene (*vipA*), and adjacent to a putative promoter with 8/12 identical nts compared to the consensus sigma70-dependent promoter (Fig. 3A). The +1 is consistent with paired-end RNAseq results we have reported prior (29). Because the majority of 5’ untranslated regions (UTRs) in *V. cholerae* are 20-40 nt, with few exceeding 300 nt (41), we speculate that the 320 nt 5’ UTR of the major T6 gene cluster may be post-transcriptionally regulated, beyond the sRNA interactions already described near the ribosome binding site (RBS) (42). Alignment of the IGRs of C6706 and V52 reveals a single SNP at -68, with a guanine (G) in C6706 at that position and a thymine (T) in V52 (Fig. 3A).

The replacement of the C6706 IGR with V52 was effectively a G-68T mutation (Fig. 3B-C), thus we further tested whether G was necessary for QstR activation by replacing the T with a G at position -68 (T-68G) in the 3223-74 WT, *qstR\**, and *ΔqstR* backgrounds. The T-68G mutation significantly increases *E. coli* survival and decreases T6 transcription in WT 3223-74 and the *ΔqstR* derivative, with killing restored in the strain with the *qstR*\* allele (Fig. 3D-E). Thus, a G at position -68 confers inducible, QstR-control, while a T results in constitutive killing *in vitro*, consistent with results recently reported (43). Based on these results we predicted this SNP is a result of adaptive evolution to control T6 activity in different environments.

### The SNP at -68 is evolutionarily conserved

To determine whether the SNP at -68 is prevalence in *V. cholerae*, we aligned the T6 IGR sequences of diverse strains that we have characterized prior for T6 killing activity (Fig. 4A) (31). Consistent with prior studies (11, 14, 16, 18), our phylogenetic analysis (Methods) of the T6 IGRs places human strains in a distinct clade, with the exception of two O1 strains isolated nearly a century ago (NCTC8457 and MAK757), and two non-O1 strains (MZO-2 O14 and V52 O37; Fig. S2). All 23 environmental isolates carry the T-68 SNP and displays constitutive T6 activity, with one exception that is chitin-inducible (1496-86) (Fig. 4A, S4). By contrast, the 18 human isolates tested carry either G or T at the -68 position (Fig. 4A, S5). The 13 chitin-inducible human isolates carry a G; five show constitutive activity and carry a T like environmental strains, with one exception that is constitutive yet carries the G (2010EL-1749) (Fig. 4A, S5). Neither C nor A are observed at -68 in any stains tested, although both pyrimidine nucleotides (T and C) confer constitutive killing at -68, and both purines (G and A) behave similarly (Fig. S3). The focal SNP location is distal from the promoter, but inconsistent with AT-rich “UP elements” that reside immediately upstream of the promoter at -38 to -59 and interact directly with the alpha subunit of RNAP (44). We propose the SNP is more likely a component of a CRE for a TF to be determined. Indeed, transversion mutations have greater effects of TF binding than transitions, as noted here (Fig. S3) likely due to changes in shape of the DNA backbone or DNA-amino acid contacts (45, 46).

**Figure 4.**
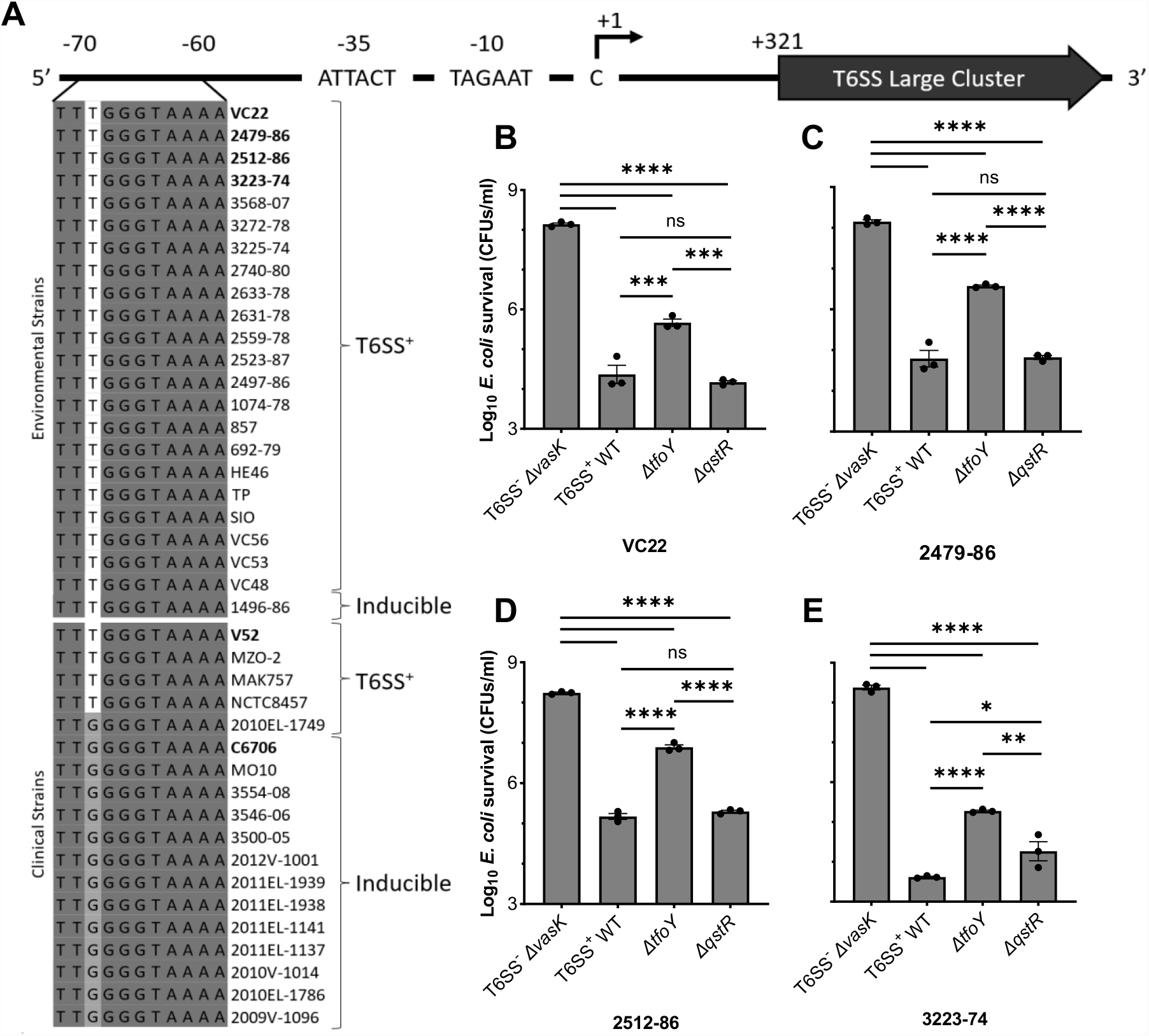
Environmental *V. cholerae* isolates encode a T at position -68 while human, chitin-induced isolates encode a G. (A) A SNP at position -68 in the IGR of the major T6 cluster controls killing activity. Conserved nts are in dark grey and the SNP of interest is highlighted in white/grey. T6 control was categorized as described (31). (B-E) Survival of *E. coli* following competition assays with WT *V. cholerae* strains and mutants was determined by CFU counts. Data shown are the mean ± S.E. of three independent experiments. A one-way ANOVA with Tukey post-hoc test was conducted to determine the significance - ns: not significant, ****p ≤ 0.0001, ***p ≤ 0.001, **p ≤ 0.01, *p ≤ 0.05.

We examined regulation of three additional genetically manipulatable environmental strains (VC22, 2479-89, and 2512-86) that exhibit T6 killing (31). Like 3223-74, QstR is expendable in each strain (Fig. 4B-E) while TfoY contributes to some extent in activating T6, with varying *E. coli* recovery observed in each derivative carrying the *ΔtfoY* mutation (Fig. 4B-E). Taken together, our findings reveal that the constitutive T6 killing activity of environmental *V. cholerae* is driven by a T at position -68, which obviates the QstR requirement, and permits modest TfoY regulation.

Bacterial adaptation to unexploited niches can be the result of horizontal gene transfer events (5) as well as mutations in protein coding and promoter regions (47, 48). Here we describe an intergenic non-coding SNP that coordinates adaptation by altering T6 control between two states – one that in inducible and the other that displays constitutive activity. While the first Type VI Secretion System was first described in *V. cholerae* in 2006, the knowledge of its regulation remains largely restricted to human isolates and incomplete, with the identity of a TF that directly controls the major T6 cluster elusive to this date (22, 24). We speculate that the focal SNP we identified at position -68 is a component of a CRE that contributes to pathoadaptation (Fig. 3A), a result of adaptive evolution, which allows *V. cholerae* to carefully control the T6SS expression in specific environments. Our results are consistent with the hypothesis that constitutive T6SS is beneficial in aquatic environments outside a human host (49), with varying degrees of TfoY contribution, which may act directly or indirectly at the transcriptional or posttranscriptional level (Fig. 3A and Fig. 4B-E, S4, S5). During human infection where selection promotes dampened T6SS, *V. cholerae* with a T-to-G mutation (inducible T6) are favored. In fact, T6-deficient human isolates (e.g. O395) have been reported to have less competitive fitness in human intestinal colonization and infection (19, 50). Although low level, basal expression of T6 contributes to pathogenesis of C6706 (51), overexpression of T6SS may be deleterious in vivo. Indeed, we have reported prior that *V. cholerae* with constitutive T6SS induces violent peristaltic contractions in a fish host (52), which may disrupt the interaction between *V. cholerae* and the gut microflora.

There remains a pressing public health need to understand the emergence of pathogens from environmental reservoirs (53). Efforts such as Microbial Genome Wide Association Studies (54) to identify genetic variants in genomes that are associated with phenotypes like virulence and antibiotic sensitivity, will be bolstered by knowledge of the ecological and evolutionary processes that promote pathogen-host association. Defining the plasticity of the regulatory circuity controlling the T6 weapon will provide insights into the role of polymorphisms in the evolution of this and other pathogens.

## Materials and Methods

### Bacterial growth conditions and plasmid constructions

All *V. cholerae* and *E. coli* (Table S1) strains were grown aerobically at 37 °C overnight in Lysogeny Broth (LB) with constant shaking or statically on LB agar. Ampicillin (100 µg/ml), kanamycin (50 µg/ml), chloramphenicol (10 µg/ml), spectinomycin (100 µg/ml), streptomycin (5 mg/ml), sucrose (20% w/v) and diaminopimelic acid (50 µg/ml) were supplemented where appropriate.

Plasmids (Table S2) used were constructed with DNA restriction nucleases (Promega – WI, USA), Gibson Assembly mix (New England Biolabs – MA, USA), and PCR amplification (Qiagen - Hilden, Germany) by PCR with Q5 polymerase (New England Biolabs – MA, USA), and primers (Table S3) generated by Eton Bioscience Inc (NC, USA) or Eurofins Genomics (KY, USA). All reagents were used according to the manufacturer’s instructions. Plasmids were confirmed by PCR and Sanger sequencing by Eton Bioscience Inc (NC, USA).

### *V. cholerae* mutant construction

All genetically engineered strains of *V. cholerae* were constructed with established allelic exchange methods using vector pKAS32 (55) and pRE118 (Addgene - Plasmid #43830). All Insertions, deletions, and mutations were confirmed by PCR and Sanger sequencing conducted by Eton Bioscience Inc (NC, USA). Primers used are in Table S3.

### Fluorescence microscopy

*V. cholerae* 3223-74 strains and chromosomal-labeled GFP *E. coli* were separately back-diluted 1:100 and incubated at 37 °C for 3 h. *V. cholerae* and *E. coli* were normalized to OD_600_ = 1 and mixed in a 1:5 ratio. A 2 μL aliquot of a mixed culture was inoculated on LB agar and allowed to dry. Cells were imaged before and after a 3 h incubation at 37 °C and 96-100% humidity using an Eclipse Ti-E Nikon (NY, USA) inverted microscope with a Perfect Focus System and camera previously described (11). The images were processed with ImageJ (34).

### Motility assay

Overnight cultures of *V. cholerae* were diluted to OD_600_ = 0.1, and 1 μL inoculated onto pre-dried LB plates with 0.3 % agar. Cells were incubated at 37 °C statically overnight, with motility determined by measuring the swarming diameter.

### Transformation assay

Chitin-induced transformation frequency was measured as described with defined artificial sea water (450 mM NaCl, 10 mM KCl, 9 mM CaCl_2_, 30 mM MgCl_2_·6H_2_O, and 16 mM MgSO_4_·7H_2_O; pH 7.8) (56). Bacteria were incubated with extracellular DNA in triplicate wells containing crab shell tabs, and transformation frequency calculated as Spectinomycin resistant (Spec^r^) CFU ml^−1^ / total CFU ml^−1^.

### QS-dependent Luciferase assay

Overnight cultures of the bacterial strains were diluted to OD_600_ = 0.001 in liquid LB in microtiter plates and incubated at 37 °C with shaking. The OD_600_ and luminescence were measured each h with a BioTek (VT, USA) Synergy H1 microplate reader to calculate Relative Luminescence Units (RLU) as Luminescence/OD_600_. Data were collected when OD_600_ = 0.6-0.8. LB medium was used as the blank for the OD_600_ and luminescence.

### GFP transcriptional reporter quantification

Overnight cultures of *V. cholerae* or *E. coli* were diluted 1:100 and incubated at 37 °C for 3 h. To enhance the translation of *gfp*, the sequence of the native RBS (12 nt sequence) was replaced with the T7 RBS (12 nt sequence) in the primers used to make the fusions. Cm was added to maintain the plasmid-borne versions of reporters that were cloned into plasmid pSLS3. 300 μL aliquots were transferred to black microtiter plates to read the OD_600_ and GFP fluorescence (Excitation: 485, Emission: 528) with a BioTek Synergy H1 microplate reader (VT, USA) to calculate Relative Luminescence Units (RLU) as Luminescence/OD_600_. LB medium was used as the blank for the OD_600_. Strain lacking reporters served as blanks for GFP fluorescence. RFU was calculated by blanked GFP fluorescence / blanked OD_600_.

### T6-mediated killing assay

Overnight cultures of *V. cholerae* or *E. coli* were back-diluted 1:100 and incubated at 37 °C for 3 h. *V. cholerae* strains and the Cm^r^ *E. coli* target were normalized to OD_600_ = 1 and then mixed at a ratio of either 10:1 or 1:5. A 50 μL mixed culture was spotted onto LB agar and dried. After a 3 h incubation at 37°C, cells were resuspended in 5 ml of LB, and serial dilutions were conducted. Finally, the resuspension was inoculated on a LB agar containing Cm to select for the surviving *E. coli*, which was incubated overnight at 37 °C and the *E. coli* colonies were counted and shown as CFU mL^-1^.

### RNA extraction and determination of the +1 of transcription by 5’-RACE

Overnight cultures of *V. cholerae* were back-diluted 1:100 and incubated at 37 °C for 3 h before lysing. Three independent cultures of T6-active *V. cholerae* C6706 *qstR\** and 3223-74 WT were harvested by centrifugation at room temperature. RNA isolation, genomic DNA removal, and RNA clean-up were performed as previously described (57). Genomic DNA contamination was confirmed by conducting PCR with primer pair specific for 16S rRNA loci (*rrsA*) as previously described (Table S3) (58). RNA purity was confirmed by NanoDrop (260 / 280 ≈ 2.0).

5’-RACE (Invitrogen™ - MA, USA) was conducted according to the manufacturer’s protocol with slight modifications. Specifically, SuperScript™ IV reverse transcriptase (Invitrogen™ - MA, USA) was used to complete the first strand cDNA synthesis. Two *vipA-*specific primers (GT3056 and GT3060) were used to identify the +1 of transcription for the major T6 gene cluster (Table S3). PCR products were purified with QIAquick PCR purification kit (Qiagen - Hilden, Germany) or Zymoclean gel DNA recovery kit (Zymo Research - CA, USA). Sanger sequencing was conducted by Eton Bioscience Inc. (NC, USA) with the corresponding nesting primer (Table S3).

### Genomic and phylogenetic analysis

Genome sequences of *V. cholerae* strains were collected from NCBI Genome database (Table S4) (59). The IGR upstream of major T6 cluster was extracted, aligned, and presented using BLAST+ v2.2.18 (60), MUSCLE v3.8 (https://www.ebi.ac.uk/Tools/msa/muscle/) (61, 62), and ESPript 3.0 (https://espript.ibcp.fr/) (37). The DNA sequence of the IGR was used for phylogenetic analysis, and the phylogenetic tree was constructed by the Maximum likelihood method using MEGA11 (63, 64).

For 2012V-1001, 2011EL-1939, 2011EL-1938, and 2011EL-1141 that do not have genome sequence available, colony PCR was conducted to amplify the 5’ IGR of the major T6 cluster using OneTaq DNA Polymerase (New England Biolabs – MA, USA). PCR products were confirmed with gel electrophoresis and Sanger sequencing by Eton Bioscience Inc. (NC, USA) with the identical primer pair (Table S3).

## Acknowledgements

We would like to thank Dr. Jyl S. Matson for assistance with RNA isolation and Dr. Marvin Whiteley and current and past members of the Hammer Lab for critiques and discussion, specifically, Dr. Samit Watve and Rakin Choudhury for bioinformatic advice and assistance.

