## Supplemental materials for "Evolution of a *cis*-acting SNP that controls Type VI Secretion in *Vibrio cholerae*"

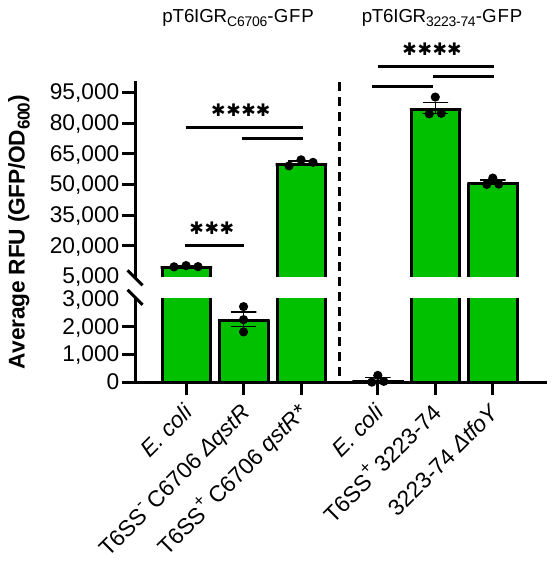

**Figure S1. The major *V*. *cholerae* T6 promoter is not constitutively expressed in *E. coli*.** *V. cholerae* or *E. coli* carrying a plasmid-encoded *gfp* gene driven by either the C6706 or 3223-74 5’ T6 IGR was grown in liquid LB with Cm. *gfp* is represented as relative fluorescent units per OD_600_ (RFU). Data shown are the mean ± S.E. from one experiment, with similar results were obtained in at least two other independent experiments. A one-way ANOVA with Tukey post-hoc test was conducted to determine the significance: ****p ≤ 0.0001, ***p ≤ 0.001.

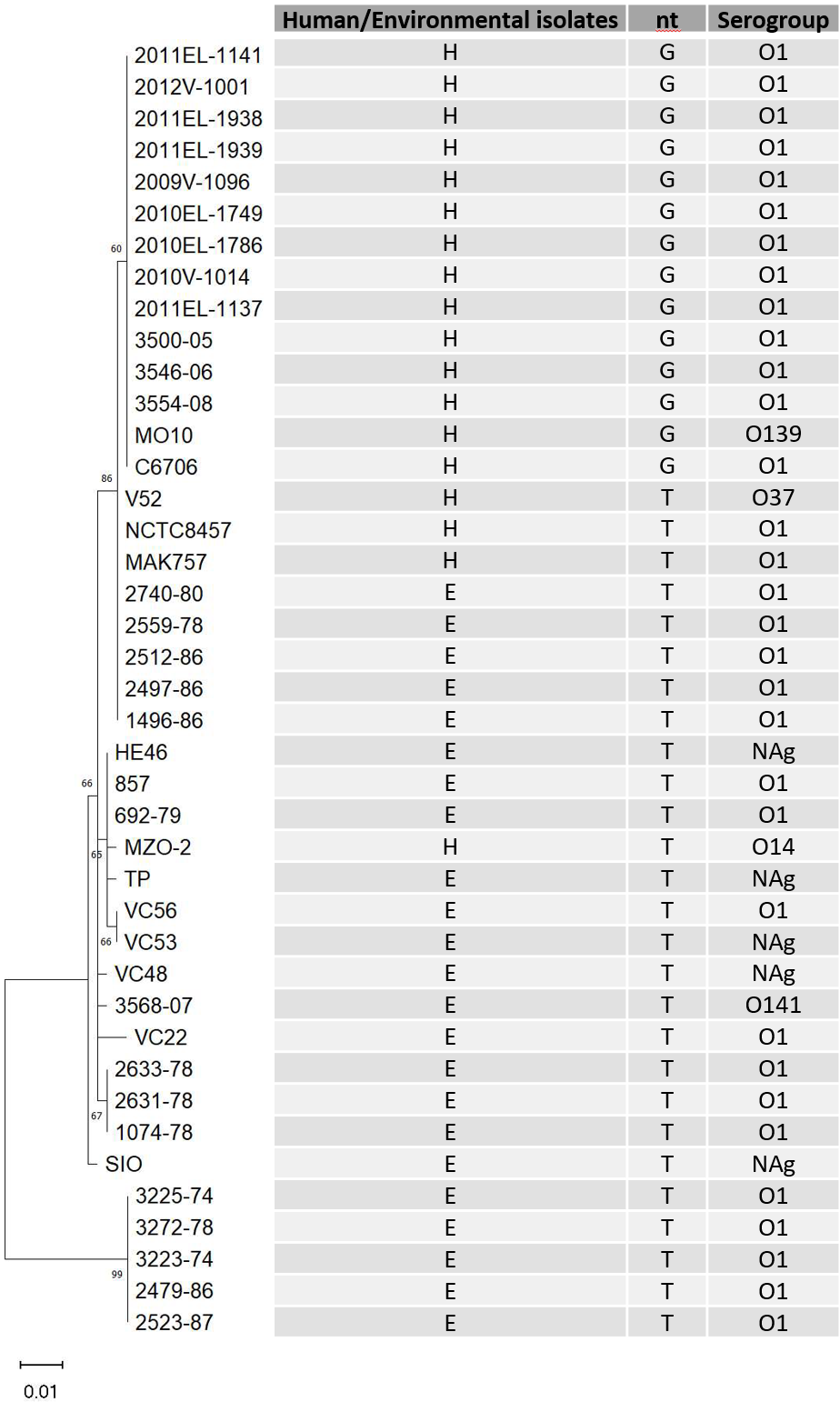

**Figure S2. Most human isolates are in a clade distinct from environmental isolates.** The T6 5’ IGR sequences of the *V. cholerae* strains described in (1) were used to conduct the maximum likelihood phylogenetic analysis with MEGA. NAg: Non-agglutinating; H: Human isolates; E: Environmental isolates.

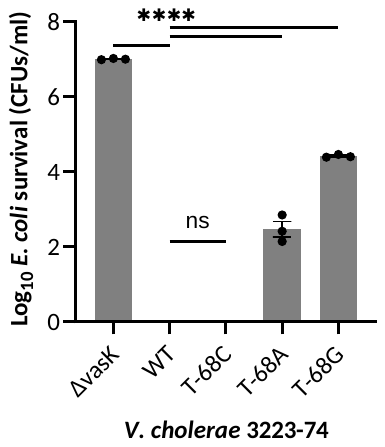

**Figure S3. Transversions at -68 alter T6 control.** Transversion mutations (T-68A, T-68G) but not a transition mutation (T-68A) introduced into the 3223-74 IGR change T6 control. Competition assays were conducted by co-culturing *V. cholerae* with Cm^R^ *E. coli* at a ratio of 1:10 for 3 h on LB agar plates. Survival *E. coli* was selected by Cm and determined by counts of colony forming units (CFUs). Data shown are the mean ± S.E. from one experiment, with similar results were obtained in at least two other independent experiments. A one-way ANOVA with Dunnett post-hoc test was conducted to determine the significance: ns: not significant, ****p ≤ 0.0001.
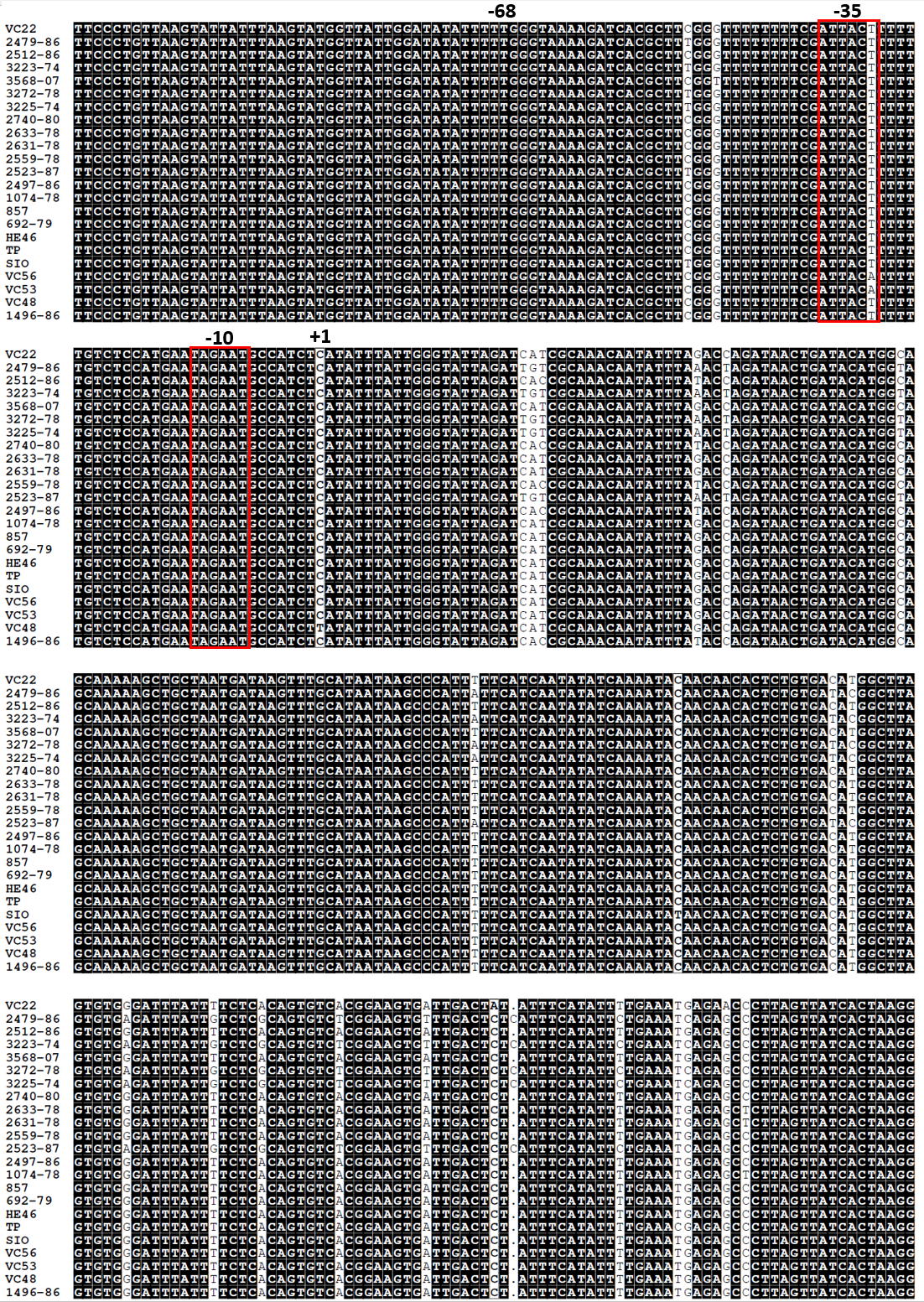

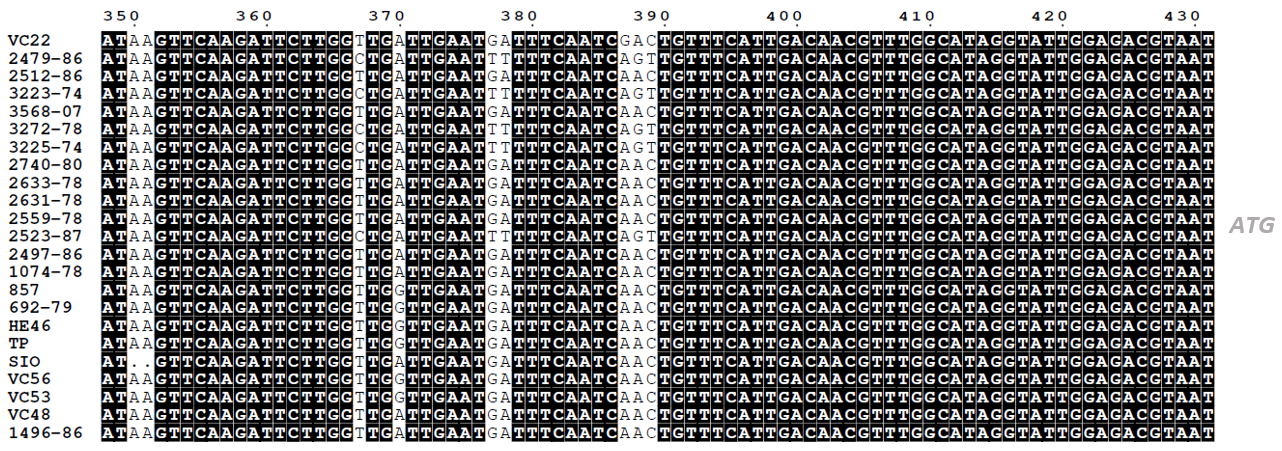

**Figure S4. Alignment of T6 IGR of environmental isolates.** Sequences of the *V. cholerae* environmental strains described in (1) were collected from NCBI database (Table S4). The T6 5’ IGR sequences were aligned using MUSCLE and generated using ESPript. Conserved bases are highlighted in black, the putative promoter is boxed, and the start codon of *vipA* is in grey.

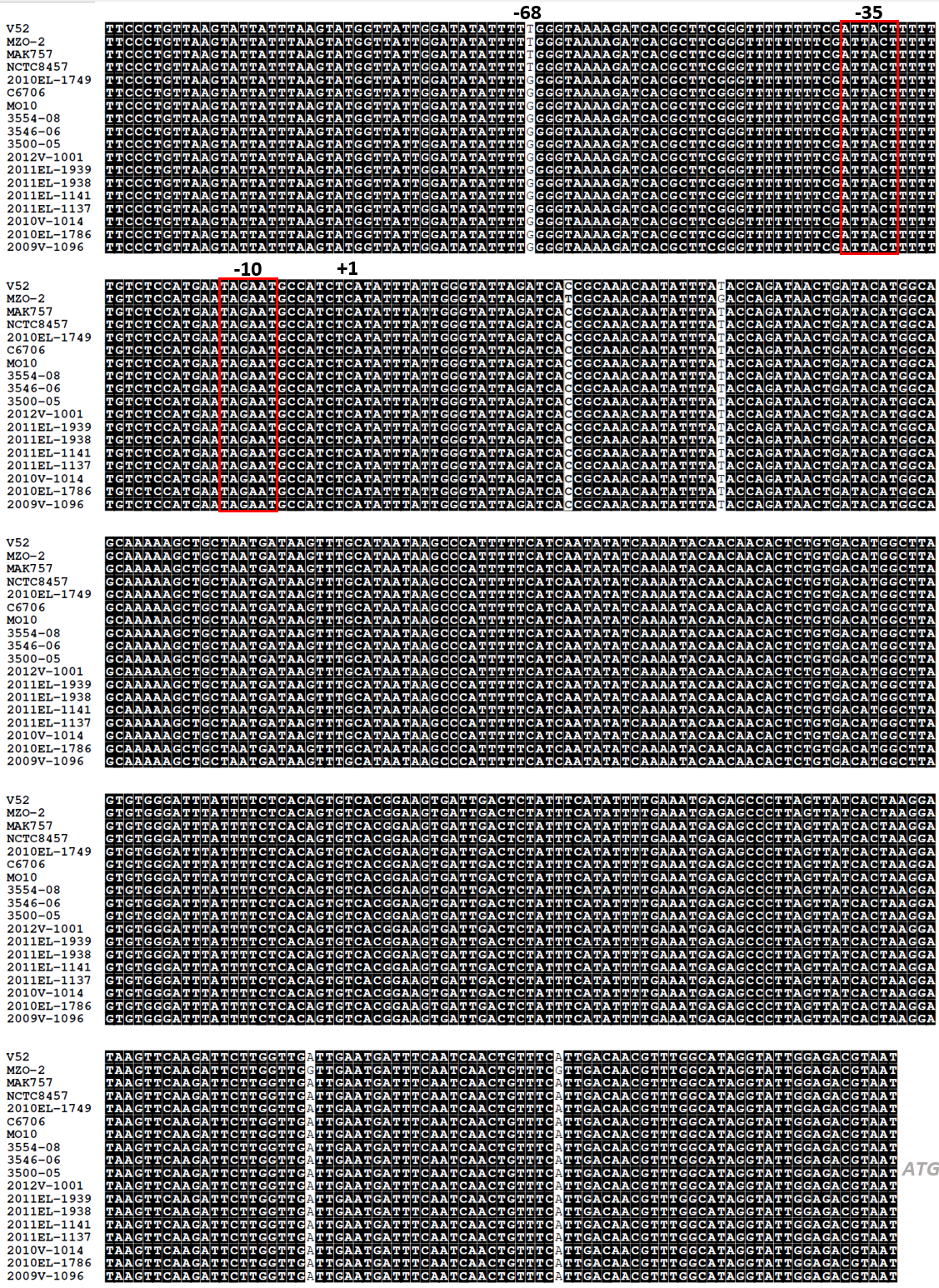

**Figure S5. Alignment of T6 IGR of human isolates.** Sequences the *V. cholerae* human-derived strains described in (1) were collected from NCBI database (Table S4), except 2012V-1001, 2011EL-1939, 2011EL-1938, and 2011EL-1141 that were generated by Sanger sequencing. The T6 5’ IGR sequences were aligned using MUSCLE and generated using ESPript. Conserved bases are highlighted in black, the putative promoter is boxed, and the start codon of *vipA* is in grey.

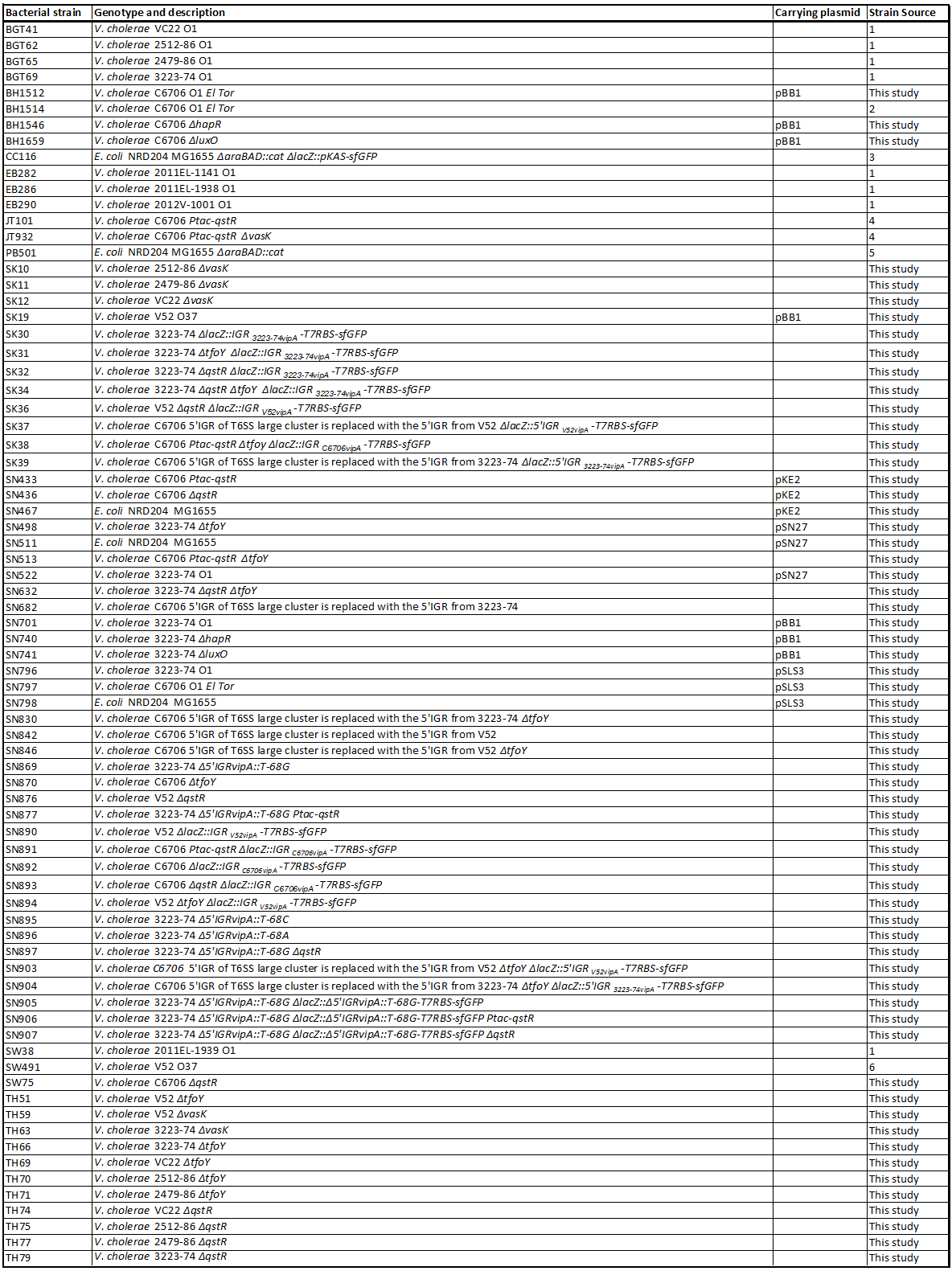

**Table S1**. **List of bacterial strains.**

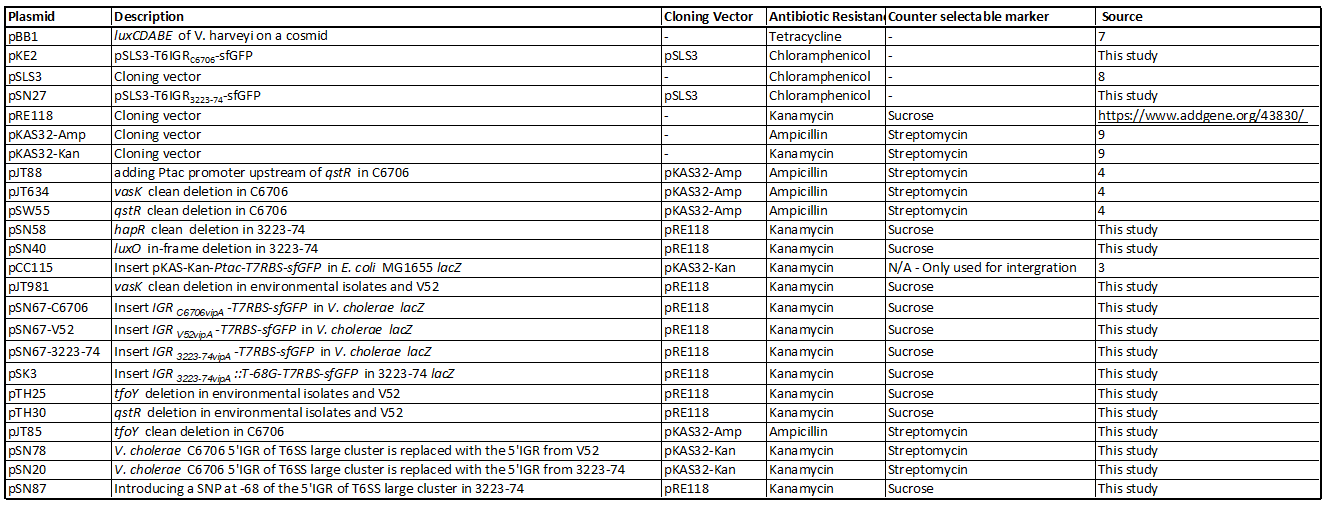
**Table S2**. **Plasmid list.**

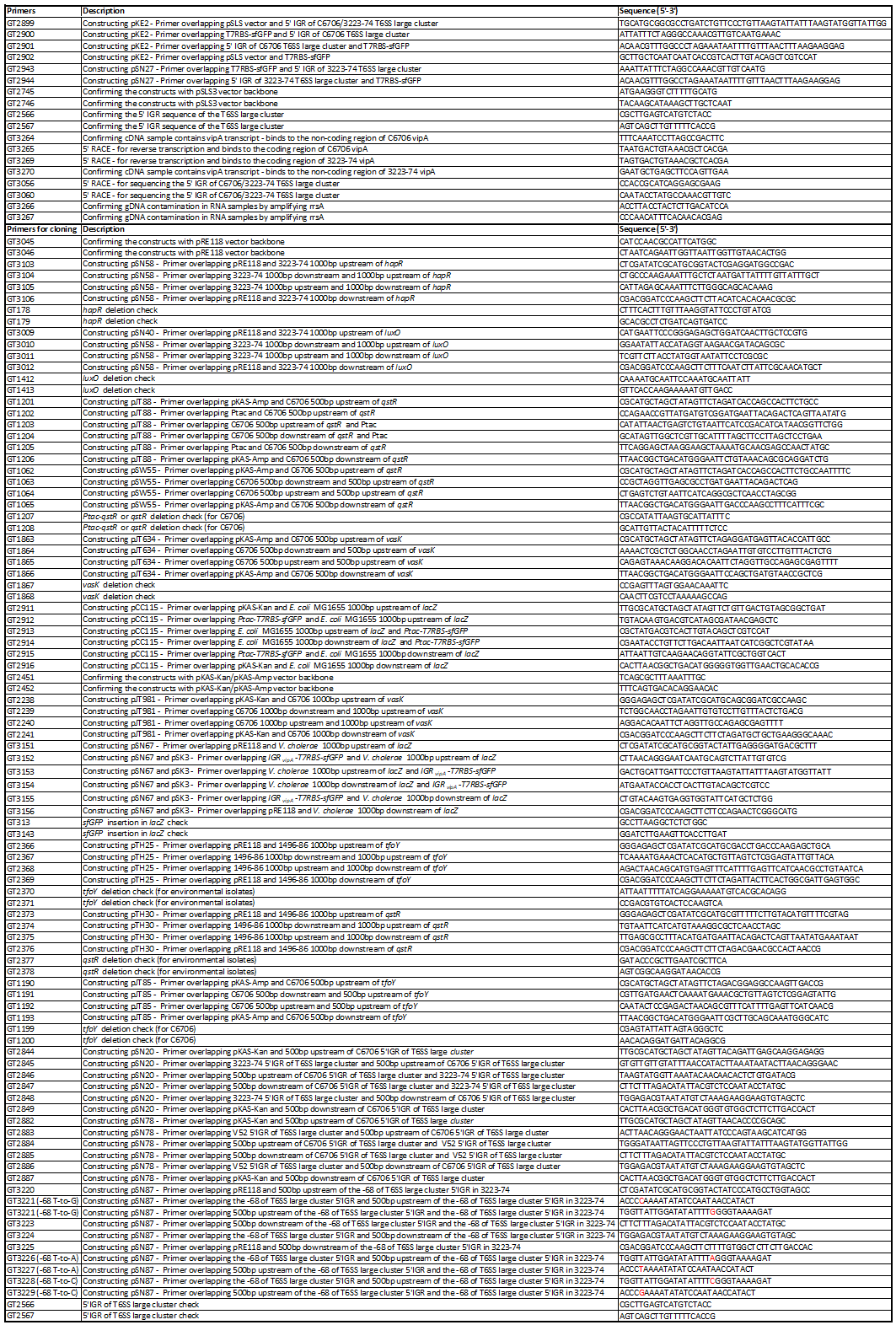

**Table S3**. **Primer list.**

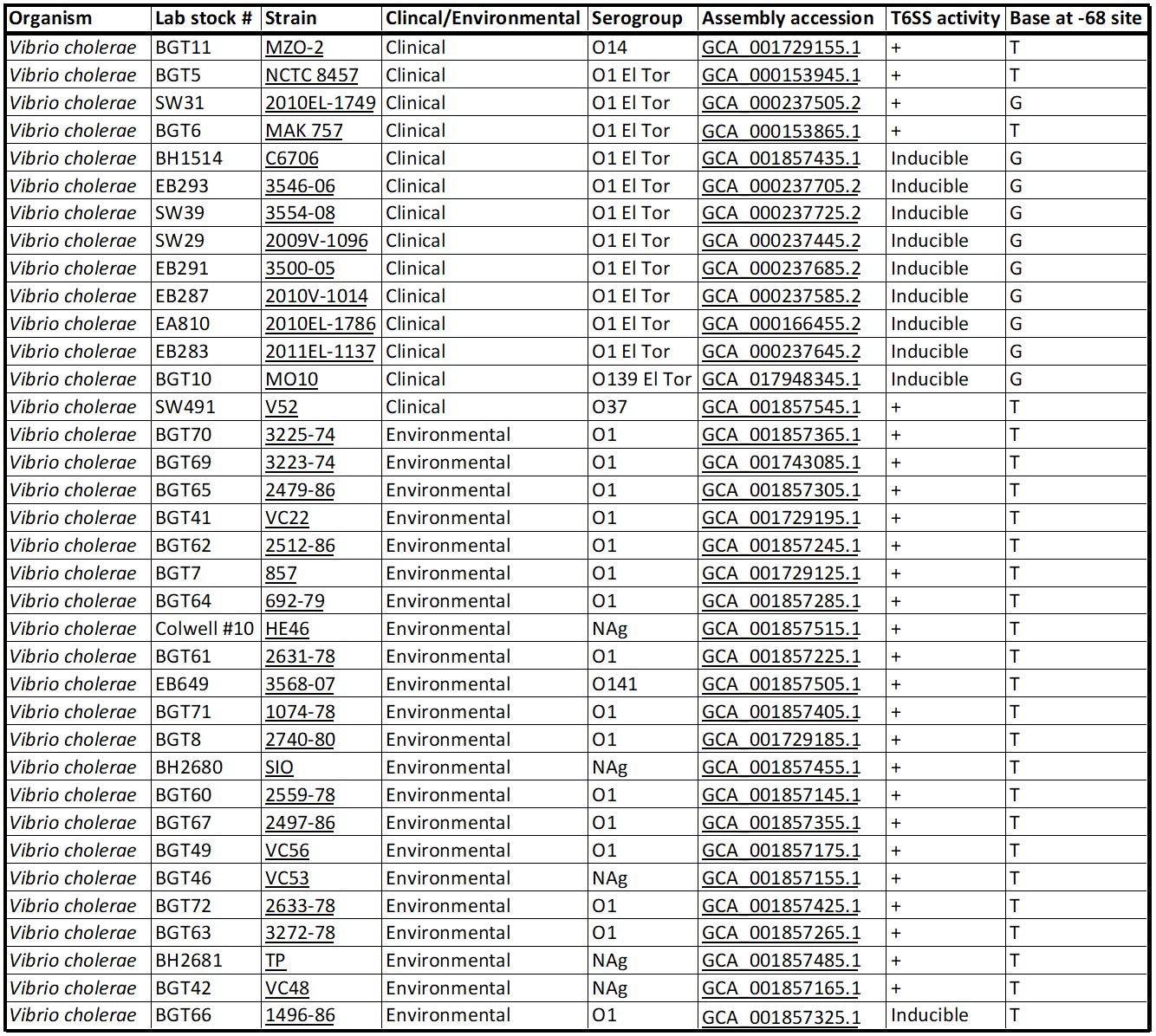

**Table S4**. **Genome list with strains details.**
